# Design of diverse, functional mitochondrial targeting sequences across eukaryotic organisms using variational autoencoder

**DOI:** 10.1101/2024.08.28.610205

**Authors:** Aashutosh Girish Boob, Shih-I Tan, Airah Zaidi, Nilmani Singh, Xueyi Xue, Shuaizhen Zhou, Teresa A. Martin, Li-Qing Chen, Huimin Zhao

## Abstract

Mitochondria play a key role in energy production and cellular metabolism, making them a promising target for metabolic engineering and disease treatment. However, despite the known influence of passenger proteins on localization efficiency, only a few protein-localization tags have been characterized for mitochondrial targeting. To address this limitation, we exploited Variational Autoencoder (VAE), an unsupervised deep learning framework, to design novel mitochondrial targeting sequences (MTSs). *In silico* analysis revealed that a high fraction of generated peptides are functional and possess features important for mitochondrial targeting. Additionally, we devised a sampling scheme to indirectly address biases arising from the differences in mitochondrial protein import machinery and characterized artificial MTSs in four eukaryotic organisms. These sequences displayed significant diversity, sharing less than 60% sequence identity with MTSs in the UniProt database. Moreover, we trained a separate VAE and employed latent space interpolation to design dual targeting sequences capable of targeting both mitochondria and chloroplasts, shedding light on their evolutionary origins. As a proof-of-concept, we demonstrate the application of these artificial MTSs in increasing titers of 3-hydroxypropionic acid through pathway compartmentalization and improving 5-aminolevulinate synthase delivery by 1.62-fold and 4.76-fold, respectively. Overall, our work not only demonstrates the potential of generative artificial intelligence in designing novel, functional mitochondrial targeting sequences but also highlights their utility in engineering mitochondria for both fundamental research and practical applications in biology.

## Introduction

A eukaryotic cell is a highly intricate entity, comprising multiple organelles responsible for various cellular processes. These organelles are enclosed in membranes and provide an optimal physicochemical environment to a multitude of nuclear-encoded proteins to carry out essential functions. Therefore, accurate localization of these proteins is crucial for maintaining cellular organization, ensuring correct metabolism, and thereby facilitating the efficient functioning of a eukaryotic cell. In nature, protein localization relies on intricate signals stored in the amino acid sequence. Over the years, significant progress has been made in understanding protein localization, aided by the characterization of localization tags in model organisms^1,2^ and the recent development of supervised machine learning (ML) models^3,4^. These advances have substantially improved our ability to target proteins to desired subcellular locations, paving the way for applications in synthetic biology^5^, chemical production^6,7^, and therapeutic interventions^8^.

The multifaceted contributions of mitochondria to cellular metabolism have made them an important target for metabolic engineering^9^ and disease treatment^10^. Mitochondria, the powerhouse of eukaryotic cells, play a pivotal role in various cellular processes, including energy production, metabolism, and apoptosis. They harbor the tricarboxylic acid (TCA) cycle and are responsible for the biosynthesis of several cofactors and metabolites. As a result, several metabolic pathways have been localized to the mitochondria, utilizing the precursor pool to boost the production of various fuels, precursor chemicals, and pharmaceuticals^9^. The mitochondrial matrix provides a distinctive physiological environment, characterized by higher pH, lower oxygen concentration, and higher reducing redox potential than the cytosol, that has proven advantageous for the expression of exogenous biocatalytic machinery^5^. Moreover, dysfunctional mitochondria, particularly those harboring inherited or acquired mutations in mitochondrial DNA (mtDNA) are associated with many diseases^11^, including Leber’s hereditary optic neuropathy (LHON), Mitochondrial Encephalopathy, Lactic Acidosis, and Stroke-like episodes (MELAS) syndrome, Leigh syndrome and others. Therefore, improving mitochondrial targeting will also aid in delivering necessary nucleases and drug molecules efficiently.

To direct proteins to mitochondria, a targeting peptide needs to be added to the N-terminus of the nuclear-encoded protein^12,13^. This positively charged, amphiphilic peptide is recognized and imported by the multi-subunit translocase of the outer and inner mitochondrial membrane, TOM and TIM complex. Upon arrival in the matrix, the peptide is cleaved by a mitochondrial processing peptidase (MPP), after which the protein folds into its mature form (Fig. 1a). However, only a few of these peptides are well-characterized, resulting in their repeated use. Given that the targeting efficiency of the peptide is dependent on the passenger protein^14^, use of a suboptimal mitochondrial targeting sequence (MTS) can lead to little to no import of the desired enzyme and result in partial compartmentalization. Furthermore, overuse of a single, endogenous targeting sequence may saturate the import machinery, resulting in potential competition with native proteins during the import system, and its downstream accumulation can compromise mitochondrial protein biogenesis and integrity^15,16^. Additionally, recurrent use of the targeting sequence to compartmentalize enzymes of a multi-gene metabolic pathway integrated into a strain can suffer from genetic instability arising from homologous recombination between sequences with high similarity. Therefore, it is highly desirable to design and characterize a toolkit of diverse, functional mitochondrial targeting sequences.

**Fig. 1:**
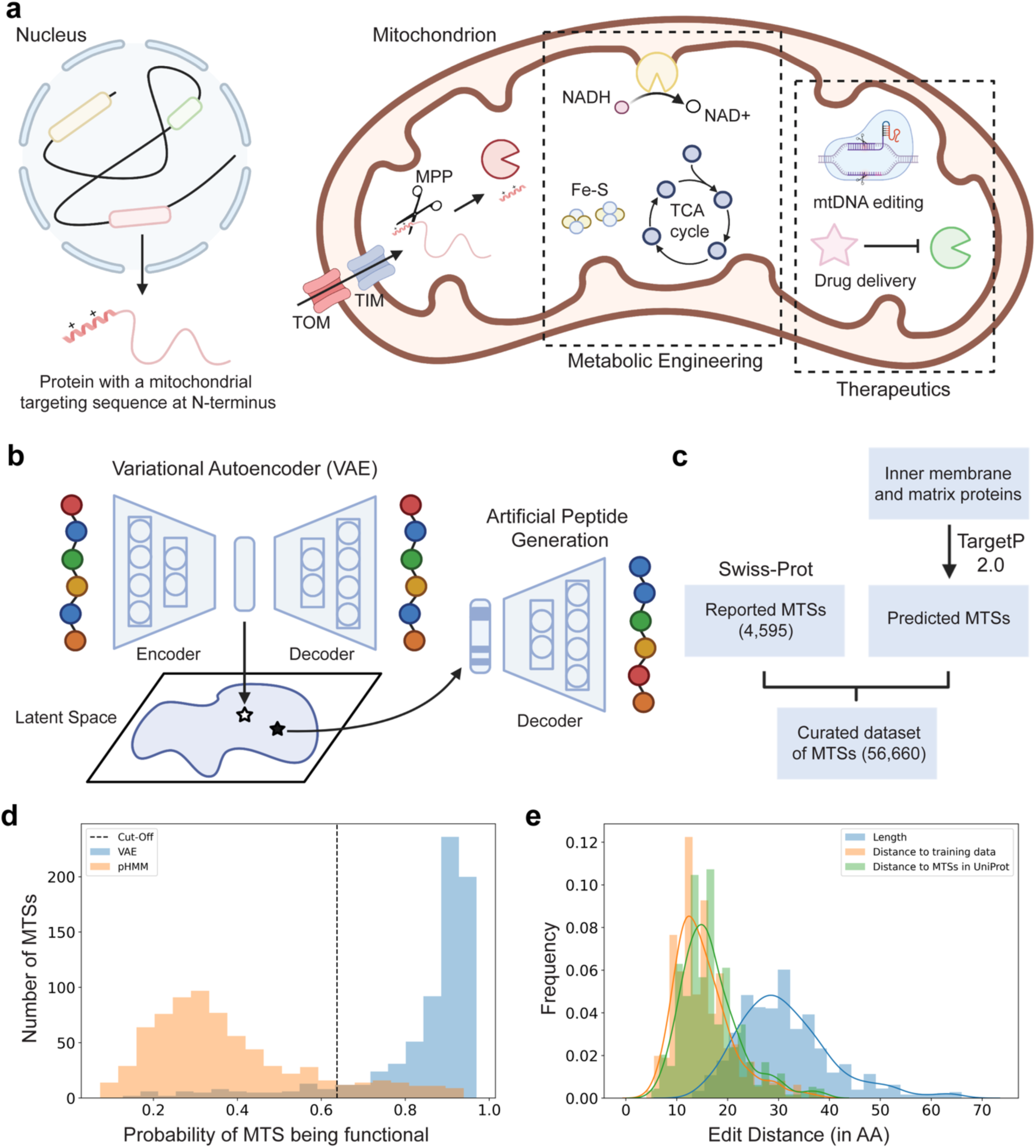
Variational Autoencoder for generation of mitochondrial targeting sequences (MTSs). **a)** Most mitochondrial proteins (99%) are nuclear-encoded and feature an N-terminal sequence for recognition and translocation by the TOM/TIM complex. Upon import into the mitochondrial matrix, the N-terminal sequence undergoes cleavage by MPP and the protein folds. Therefore, utilizing such a targeting sequence enables the delivery of enzymes and drugs for diverse applications, including biochemical production, mtDNA editing, and treating mitochondrial disorders. **b)** Scheme for generating artificial MTSs using Variational Autoencoder (VAE). The model receives a one-hot encoded representation of MTS as an input. The encoder compresses the input into a latent vector, and the decoder reconstructs the original input. Once the model is trained, one can feed a vector sampled from the latent space to the decoder and generate novel MTSs. **c**) Curated dataset of MTSs for training the VAE. The dataset includes MTSs reported in Swiss-Prot and TargetP 2.0-predicted MTSs for proteins with mitochondrial matrix or inner membrane as subcellular location. **d)** Predicting functionality of generated MTSs using DeepLoc 2.0. A set of 730 VAE and pHMM-generated MTSs, appended with GFP, were analyzed for their ability to target mitochondrion. Out of 730, 658 sequences (90.14%) were deemed functional for VAE compared to 89 sequences (12.2%) generated using pHMM. **e)** VAE-generated MTSs are diverse in sequence. Many of the generated sequences are 10 to 15 mutations away from the MTSs in the training data and UniProt. Note that the percentage of AMTS-tagged GFP localized to mitochondrion was calculated based on the probability threshold of 0.6373 specified on the DeepLoc 2.0 server. TOM/TIM: translocase of the outer/inner membrane; MPP: mitochondrial processing peptidase; pHMM: profile Hidden Markov Model.

In recent years, ML approaches have shown tremendous promise in peptide design. Specifically, deep generative models have demonstrated their ability to capture intricate patterns solely from sequence data^17^. By learning from vast datasets of unlabeled peptides, these models can generate novel sequences that adhere to the underlying rules governing peptide function and structure. Therefore, various model architectures are employed to design antimicrobial peptides^18^, signal peptides^19^, and cell-penetrating peptides^20^. In this study, we trained a Variational Autoencoder (VAE) to generate new-to-nature mitochondrial targeting sequences. We performed experiments to validate the functionality of these sequences in four eukaryotic organisms, including *Saccharomyces cerevisiae*, *Rhodotorula toruloides*, *Nicotiana benthamiana*, and the HEK293 cell line. Furthermore, we harnessed the latent space interpolation capabilities of the VAE to create putative sequences capable of targeting mitochondria and chloroplasts simultaneously. Through an extensive analysis of the generated sequences, we extracted insights to inform evolutionary hypotheses regarding dual targeting sequences. Finally, we showcased the practical applications of these artificially designed sequences, highlighting their effectiveness in pathway compartmentalization and enhancing our ability to target mitochondria.

## Results

### Creation of a Variational Autoencoder for designing artificial mitochondrial targeting sequences

Mitochondrial targeting sequences (MTSs) are positively charged peptides with a tendency to form amphiphilic α-helices^12,13^. These peptides are 10-120 amino acids long (with a typical length of approximately 35 amino acids) and are located at the N-terminus of the proteins destined for mitochondria. Unlike organelles such as peroxisomes, these targeting sequences do not exhibit specific motifs and instead rely on physicochemical and structural characteristics for their recognition and import into mitochondria^13^. This makes the design of a diverse peptide library challenging, considering the extensive design space (20^35^ for an MTS of common length), the lack of an evident consensus sequence, and the unpredictable relation with the passenger protein. Deep generative models are capable of learning complex, non-linear mappings and representing the underlying data distribution, enabling the design of novel peptides^17^ and proteins^21^ with desired functions from sequence information alone. In this work, we sought to utilize VAE to generate new-to-nature MTSs (Fig. 1b). VAE consists of two components: the encoder which converts the input sequence into a latent space representation, and the decoder, that reconstructs these latent representations back into the input sequence. The architecture takes the form of a bottleneck, enabling the model to capture essential features. In addition to optimizing the reconstruction loss during training, VAE minimizes the Kullback-Leibler (KL) divergence loss, effectively shaping the distribution of the latent space to fit a multivariate Gaussian distribution. By leveraging the trained model, one can sample latent vectors from the continuous space and input them into the decoder and design novel sequences that closely resemble the properties of the training data.

To facilitate effective training of VAE and encompass a comprehensive design space, we curated a dataset of MTSs. First, we utilized transit peptide annotation and mitochondria as the subcellular location to procure 4,984 MTSs from Swiss-Prot^22^, out of which 4,595 are unique. These sequences are either experimentally validated or predicted using sequence similarity and software such as Mitofates^23^, Predotar^24^, and TargetP^25^. However, this dataset is relatively small and exhibits a notable bias towards model organisms (Supplementary Fig. 1). To address this limitation and enhance dataset diversity, we leveraged TargetP 2.0^3^, a state-of-the-art attention-based deep learning model, to predict MTSs for proteins in Swiss-Prot and TrEMBL with subcellular localization specified as mitochondrial matrix and inner membrane^22^. TargetP 2.0 predicts signal or transit peptides separately for plant and non-plant organisms. Therefore, we used taxonomy classification from UniProt to segregate the resulting protein sequence dataset and fed it to TargetP 2.0 to extract MTSs. Subsequently, we refined the MTSs by filtering for peptides with valid amino acid sequences, restricting lengths to 11-69 amino acids, and eliminating duplicates. Finally, we obtained a final dataset comprising 56,660 peptides for model development (Fig. 1c).

To train the VAE, we introduced a ‘$’ symbol at the C-terminus of the peptides, marking the cleavage site. Subsequently, considering the skewed distribution of peptide length, we performed a stratified 9:1 split of the dataset, ensuring a balanced representation of peptides in both the training and validation sets (Supplementary Fig. 2). We padded all peptides to a uniform size of 70 and employed one-hot encoding to create an input representation for the encoder. Initially, we trained a recurrent neural network (RNN)-based VAE^26^ that was previously used to generate antimicrobial peptides^18^. However, post training, we observed that the same sequence was generated from different randomly initialized latent vectors, with the sequences displaying repeated amino acid residues and minimal mutations, indicating low diversity (Supplementary Table 1). We then implemented the encoder and decoder of the VAE with fully connected layers and did not observe this issue. Therefore, we selected this model for further validation. More details of the model implementation are provided in Materials and Methods.

### *In silico* analysis reveals generated MTSs are functional, highly diverse, and not found in nature

Given the trained VAE model, we generated artificial peptides and conducted a comprehensive analysis of their functionality, diversity, physicochemical attributes, and structural properties. We sampled 1000 vectors from a normal distribution N(*μ* = 0, σ = 1) and fed them to the decoder to generate new-to-nature MTSs. Subsequently, we refined these to obtain 730 peptides with valid amino acid sequences. To verify if the peptides can localize proteins to the mitochondrion, we prepended them to the Green Fluorescent Protein (GFP) lacking the start methionine and analyzed them using DeepLoc 2.0^4^, a deep learning model for predicting the subcellular localization of proteins (Supplementary Fig. 3). We observed that 90.14% of the peptides were predicted to target mitochondria (Fig. 1d). We benchmarked these sequences with an equal number of MTSs designed using the profile Hidden Markov Model (pHMM). A pHMM captures position-specific residue probabilities and insertion and deletion states, making it valuable in generating new sequences that adhere to the language of a protein family^27^. However, when prepended to GFP, only 12.2% of the artificial peptides were predicted to be functional by DeepLoc 2.0. Next, we compared the VAE-generated MTSs with the sequences in the training data and the MTSs reported in UniProt, as the latter are accessible to researchers. We calculated the Levenshtein distance to determine the number of mutations between the generated sequence and its closest match. Our analysis revealed that VAE-generated MTSs are highly diverse, averaging 10-15 amino acids away from any naturally occurring MTSs (Fig. 1e).

Next, we evaluated the model’s capability to cover the natural MTS space. We generated 1,000,000 artificial MTSs and selected peptides with valid amino sequences. Subsequently, we applied CD-HIT^28^ on the training data and VAE-generated MTSs and obtained natural and artificial MTSs with less than 30% sequence identity, respectively. Next, we embedded them with UniRep model^29^ and employed Uniform Manifold Approximation and Projection (UMAP) to visualize the latent space in two dimensions. This visualization revealed three distinct clusters, with artificial MTSs effectively covering the entire natural sequence space (Fig. 2a). Upon closer examination, we noticed one cluster contained MTSs without the start methionine, a discrepancy propagated through MTSs of proteins from UniProt used for model training. While we inspected the peptides in the remaining two clusters, we failed to detect any apparent differences.

**Fig. 2:**
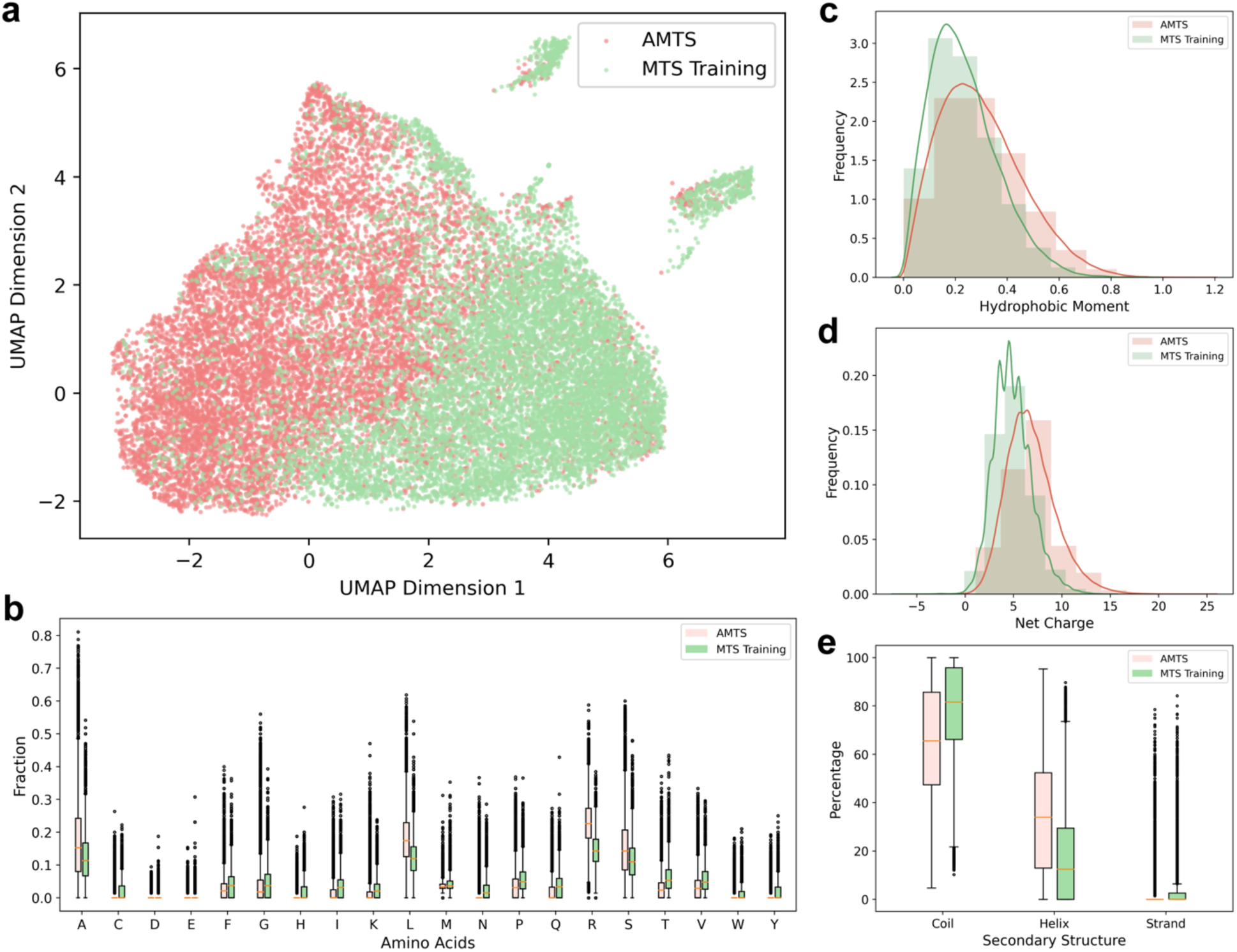
Characteristics of artificial mitochondrial targeting sequences (AMTSs). **a)** Uniform Manifold Approximation and Projection (UMAP) visualization of UniRep embeddings depicting the distribution of AMTSs and MTSs within the training dataset. VAE effectively generates AMTSs across both high and low-density regions in sequence space. Comparative analysis of physicochemical and structural attributes between generated AMTSs and MTSs in the training data, showcasing **b)** Amino acid composition, **c)** Hydrophobic moment, **d)** Net charge, and **e)** Secondary structure. Overall, AMTSs exhibit a net positive charge and form amphiphilic α-helix structures, features crucial for directing protein to mitochondria.

Furthermore, we compared amino acid composition and physicochemical characteristics of artificial and naturally occurring MTSs, as shown in Fig. 2b-d and Supplementary Fig. 4. We used BioPython to compute these features. Overall, VAE-generated MTSs followed a similar distribution for amino acid composition compared to the MTSs in the training dataset. Artificial MTSs were enriched in Alanine (A), Arginine I, Leucine (L), and Serine (S). Negatively charged residues, Aspartic (D) and Glutamic acid I, were depleted for sequences in both datasets. Therefore, artificial MTSs carried a net positive charge, an important feature for functional MTSs. Next, we calculated the mean hydrophobic moment using modlAMP^30^ to measure the amphiphilicity of the peptides^31^ and observed a slight shift towards higher values. Hydrophobicity, as measured using the Eisenberg scale, exhibited a more negative trend, while when quantified through GRAVY (Grand average of hydropathicity index), followed a similar distribution compared to the training dataset (Supplementary Fig. 4a, b). Generated MTSs also spanned the length of the peptides in the training dataset (Supplementary Fig. 4c). Next, we calculated the fraction of α-helix, β-sheet, and loop (or coil) using S4PRED^32^, a state-of-the-art model for single sequence-based secondary structure prediction. We observed artificial MTSs displayed a higher tendency to form α-helix at the expense of coil (Fig. 2e). The results demonstrate that the VAE effectively captured meaningful features essential for mitochondrial localization.

### Sampling algorithm to select sequences for *in vivo* experimental validation

Based on the confidence gained from the *in silico* analysis, we randomly selected nine VAE-generated MTSs with TargetP 2.0 scores above 0.9 for *in vivo* evaluation in *S. cerevisiae*. Using Gibson assembly^33^, we fused these sequences to the N-terminus of GFP. Additionally, we constructed a positive control using the MTS of COX4^34^, a well-characterized endogenous peptide commonly used for mitochondrial targeting. We transformed these GFP plasmids into *S. cerevisiae* and verified the mitochondrial targeting capability of the sequences through confocal microscopy. We initially examined the negative control (GFP) and the positive control (COX4-GFP). The overlap of GFP fluorescence with the mitochondrial stain, MitoTracker^TM^ Orange CMTMRos, confirmed that COX4 can localize the protein to mitochondria, while GFP on its own cannot (Supplementary Fig. 5). Similarly, we conducted the characterization for nine artificial MTSs (AMTSs). Out of these sequences, AMTS 131, 205, 225, and 335 demonstrated selective targeting to mitochondria, while AMTS 6, 64, and 96 displayed targeting to multiple subcellular locations, including mitochondria.

Next, we sought to demonstrate the functionality of these VAE-generated sequences in three eukaryotic organisms: *Homo sapiens*, *N. benthamiana*, and *R. toruloides*. While some of the artificial peptides demonstrated successful targeting, we wondered whether the interaction with the import machinery of an organism configured a bias in the amino acid sequence of MTSs. Our hypothesis also stemmed from the consideration that tools for predicting subcellular localization incorporated taxonomy in their predictions^3,23^. Therefore, we analyzed various attributes of naturally occurring MTSs in these organisms (Supplementary Fig. 6) and formulated a sampling scheme to choose sequences better suited for validation in the organism of interest (Fig. 3a). We acquired these sequences from UniProt or predicted them using TargetP 2.0. Subsequently, we refined them and calculated amino acid composition, physicochemical attributes, and secondary structure as previously described. MTSs from different organisms preferred varying amino acid residues. For instance, MTSs from the human proteome exhibited enrichment in Glycine (G) and Alanine (A), accompanied by a reduction in Serine (S) when compared to MTSs from other three proteomes. Similarly, these MTSs differed in hydrophobic moment and had a higher tendency to form an α-helix compared to fungal MTSs. Next, we used the pre-trained UniRep model to obtain embeddings for the targeting sequences, capturing amino-acid, physicochemical, and structural attributes through meaningful protein representation^29^, and visualized them in a two-dimensional space using UMAP. Based on the density plot, it was evident that the MTSs for different organisms were clustered to some extent (Fig. 3a).

**Fig. 3:**
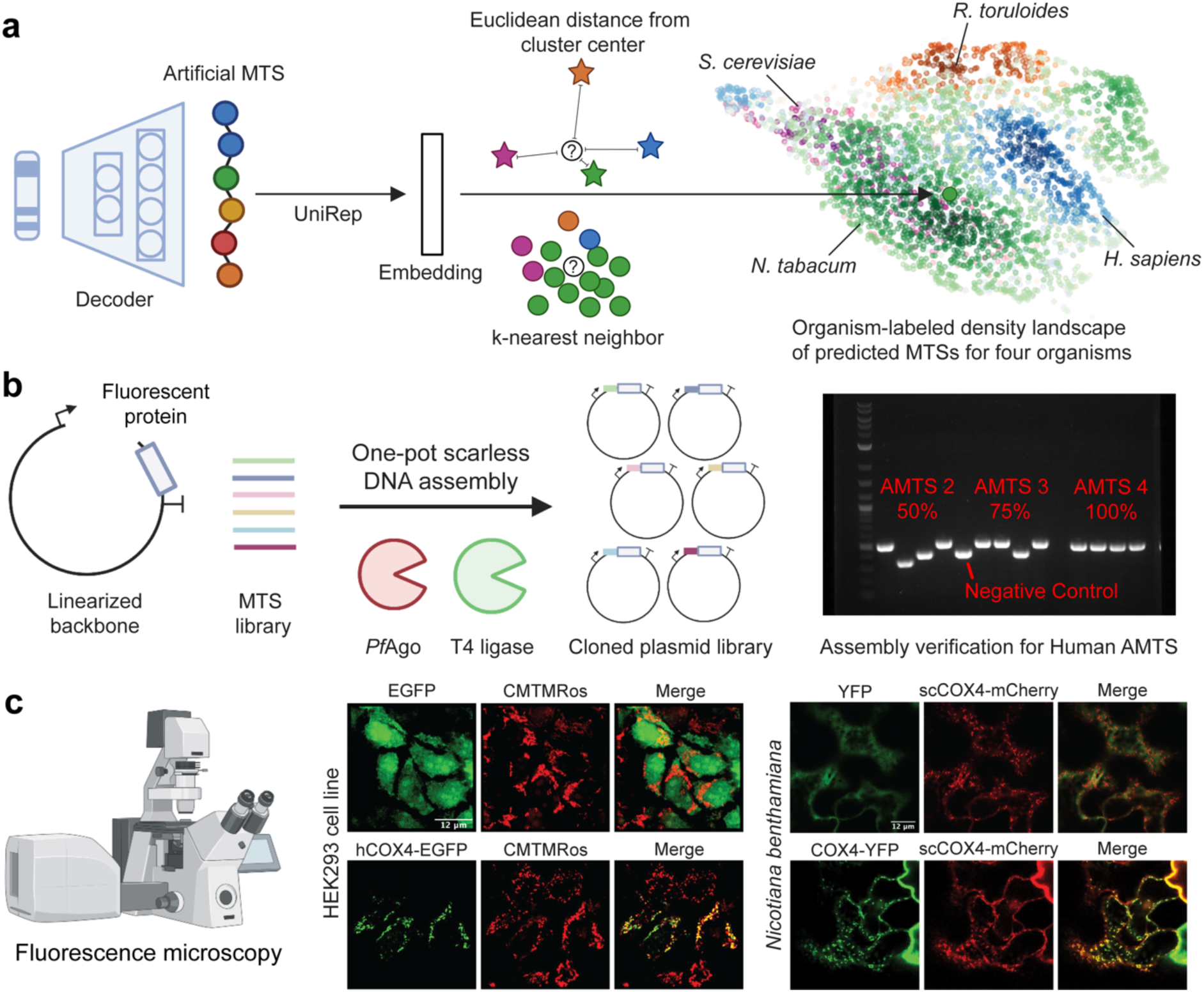
Organism labeling and workflow for *in vivo* evaluation of generated artificial mitochondrial targeting sequences (AMTSs). **a)** Selection of AMTSs for experimental characterization. AMTSs are generated through the VAE, encoded via the pre-trained UniRep model, and annotated with an organism label based on proximity to the cluster center, defined as the mean UniRep embedding of MTSs in a proteome, and using k-Nearest Neighbors. **b)** Scarless DNA assembly of AMTSs utilizing *Pf*Ago/AREs. Plasmids are cleaved using *Pf*Ago and phosphorylated guides, creating sticky ends. Annealed DNA sequences, encoding for AMTSs, are then ligated upstream of fluorescent reporter proteins using T4 DNA ligase. Final constructs are verified through Colony PCR, demonstrating high assembly fidelity. **c)** Functional validation of positive control MTSs. Plasmids harboring MTS-fluorescent protein constructs are transformed into the host organism, followed by verification of mitochondrial localization using fluorescence microscopy. Mitochondria in HEK293 cell lines are visualized with MitoTracker^TM^ Orange CMTMRos, whereas mitochondrial targeting in *N. benthamiana* is confirmed through protein colocalization with scCOX4-mCherry. Scale bar: 12 µm.

To determine the organism for characterizing an MTS, we devised a sampling scheme, implementing two criteria using the 1900-dimensional UniRep representation of MTSs. The first criterion categorizes MTSs based on their proximity to the cluster center, calculated as the average UniRep representation of MTSs associated with a specific organism. The second criterion follows the principle of k-Nearest Neighbors, assigning an MTS to an organism by computing the frequency of organism labels for its 20 closest naturally occurring counterparts. We choose an MTS for validation only when both criteria concur. To highlight the effectiveness of this sampling scheme over random selection, we analyzed the cleavage patterns of peptides in the human proteome (Supplementary Fig. 7). The consensus for the eight residues at the C-terminus revealed that the sampling results in a pattern more closely resembling naturally occurring MTSs, as anticipated. Moreover, Glycine (G), an amino acid residue favored by MTSs in the human proteome, is identified as a positive attribute only for sampled MTSs and not for the ones generated randomly, demonstrating a method for capturing organism-specific bias. Therefore, utilizing this approach, we labeled the VAE-generated MTSs and selected eight peptides based on the distance from the cluster center for characterization in four eukaryotic organisms (Supplementary Table 2).

### VAE-generated peptides target mitochondria *in vivo* across various eukaryotic organisms

To test the 32 VAE-generated sequences obtained after sampling, we fused these artificial peptides to the N-terminus of various reporter genes. Instead of using Gibson assembly, we adopted *Pf*Ago-based assembly^35^, a more versatile cloning approach. *Pf*Ago is an artificial restriction enzyme that uses single-stranded DNA guides to cleave double-stranded DNA, making sticky ends of the desired length. Unlike the previous approach, we modified the script to design guides at the end of the amplified plasmid backbone to create a 12-bp overhang. We then cloned artificial MTSs and appropriate positive controls using T4 DNA ligase. This modified approach aided in constructing plasmids with high fidelity and ease (Fig. 3b). Subsequently, we performed confocal microscopy and confirmed the negative (EGFP/YFP) and positive controls for mitochondrial localization (Fig. 3c). We provide a summary of artificial MTS validation in the four eukaryotic organisms in Fig. 4a. All eight peptides localized GFP to the mitochondria of HEK293T cells, as demonstrated by overlap with MitoTracker^TM^ Orange CMTMRos (Fig. 4b). We also verified successful peptide cleavage using Western blot (Supplementary Fig. 8). Similarly, six out of eight peptides successfully targeted Yellow Fluorescent Protein (YFP) to *N. benthamiana*’s mitochondria (Supplementary Fig. 9). We confirmed this using agrobacterium-mediated transient expression of fluorescent proteins and subsequent colocalization with ScCOX4-mCherry.

**Fig. 4:**
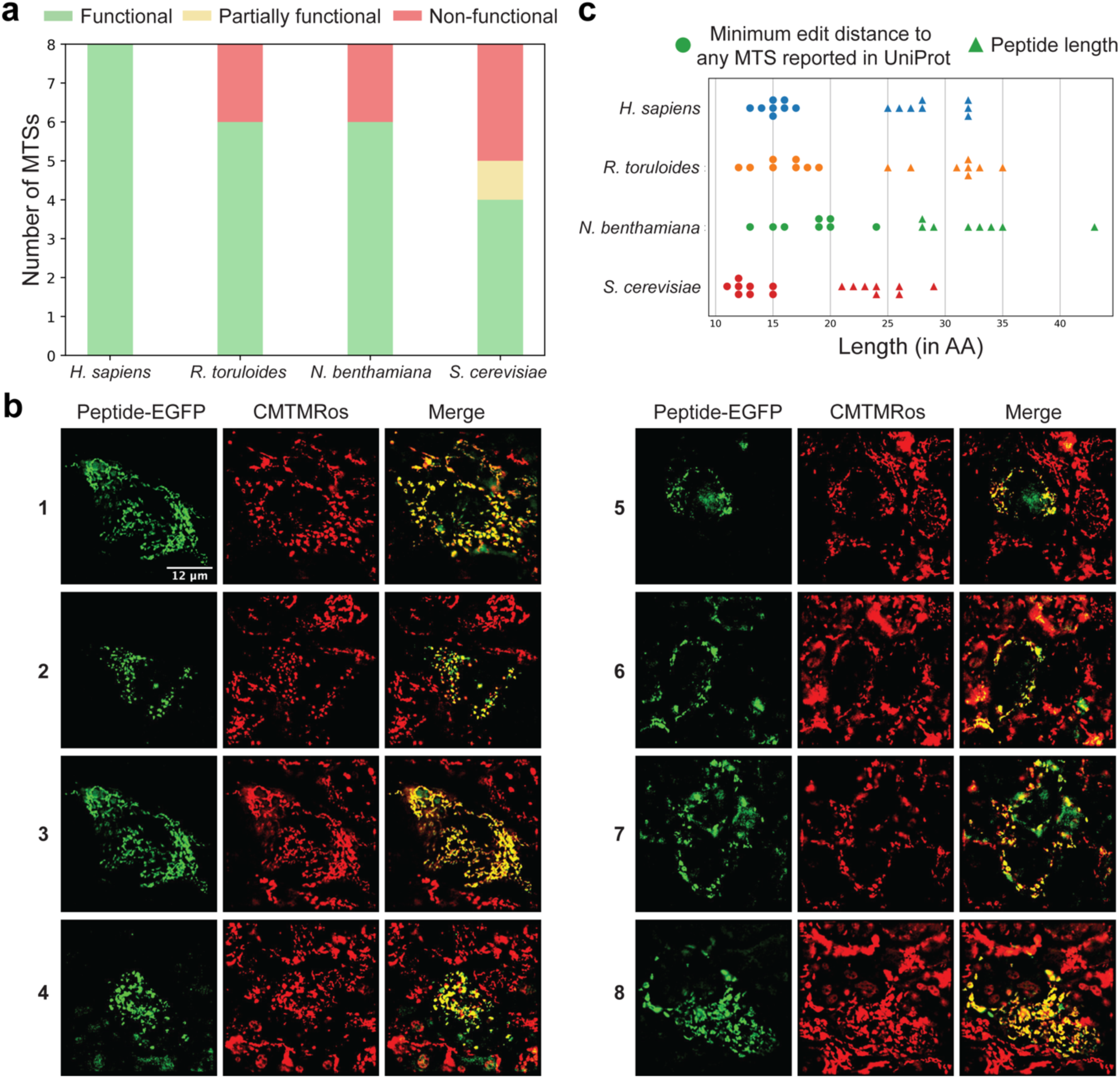
Characterization of artificial mitochondrial targeting sequences (AMTSs) in four eukaryotic organisms. **a)** Summary of AMTS functionality in HEK293 cell line, *R. toruloides*, *N. benthamiana*, and *S. cerevisiae*. **b)** Fluorescence microscopy analysis of AMTS-EGFP constructs in the HEK293 cell line. All eight sequences target mitochondria *in vivo*, confirmed through overlap with MitoTracker^TM^ Orange CMTMRos. **c)** Levenshtein distance of artificial peptides to MTSs reported in UniProt, illustrating the diversity of characterized sequences. Scale bar: 12 µm.

Next, we verified the VAE-generated MTSs in *R. toruloides*, a non-model oleaginous yeast. It is considered a promising chassis to produce chemicals and biofuels, owing to its ability to grow on diverse substrates and its high endogenous flux towards lipids and carotenoids^36^. Therefore, to characterize artificial MTSs in this host, we knocked out *crt*YB, the enzyme responsible for producing β-carotene, using a previously constructed sgRNA cassette^37^ to avoid interference with analysis. Next, we transformed the linear AMTS-tagged GFP fragment with the necessary selection marker in the strain and analyzed them using confocal microscopy. Six out of eight peptides successfully localized GFP to the mitochondria, confirmed using Rhodamine B, hexyl ester (Supplementary Fig. 10). We used TargetP 2.0-predicted MTS of RtCOX4 as a control. However, we did not observe GFP localized to mitochondria. Finally, we characterized eight more artificial MTSs in *S. cerevisiae*. Among these, four constructs showed successful mitochondrial targeting (Supplementary Fig. 11).

Overall, we achieved a success rate of 75-100% for AMTSs characterized in the HEK293 cells, *N. benthamiana*, and *R. toruloides*. However, the hit rate dropped to 50% for AMTSs tested in *S. cerevisiae*. This decrease can be partially attributed to the dispersion of naturally occurring MTSs in *S. cerevisiae* across the entire UMAP space, resulting in inefficient sampling. Notably, the characterized peptides maintained diversity even after sampling closer to the cluster center, with each peptide being 15-20 mutations away from any MTSs reported in UniProt (Fig. 4c).

### Latent space interpolation enables to design peptides capable of targeting multiple subcellular locations for dual organelle engineering

Our earlier analysis indicated that 27 and 17 of the VAE-generated peptides localize GFP to the chloroplast (plastid) or both the chloroplast (plastid) and mitochondrion (Supplementary Fig. 3), respectively. Initially, we suspected the model learned these attributes from peptides of proteins with mislabeled subcellular localization annotations and potential false positives from TargetP 2.0 predictions. However, MTS and chloroplast targeting sequences (CTSs) are evolutionarily related and possess similarities^38^. In nature, 5% of the proteins in the endosymbiotic organelles of plant cells are expected to be dual localized^39,40^. Dual localization can occur either through a protein carrying different signals or a single ambiguous signal for both organelles. Previous studies report that the characteristics of these ambiguous signals are intermediate to those of MTSs and CTSs^41^. Therefore, we asked if interpolation in learned latent space can generate peptides capable of targeting both organelles and provide insights into their evolutionary trajectories. We trained a variational autoencoder (Dual-VAE) on MTSs and CTSs from the Viridiplantae kingdom reported in UniProt or predicted using TargetP 2.0 (Supplementary Fig. 12). We observed two posterior distributions in the latent as the model learns from sequences belonging to two different classes (Supplementary Fig. 13a). Next, we utilized K-Means to locate the center of each distribution to select latent vectors for interpolation. Employing a Euclidean distance threshold of 0.4 from the cluster center, we obtained over 50 MTSs and CTSs. For each sequence pair, we performed linear interpolation and generated three artificial sequences by feeding the approximated latent vectors to the decoder (Fig. 5a). First, we analyzed these sequences for functionality using DeepLoc 2.0^4^ to predict if the sequences generated at the interface of two distributions can target both endosymbiotic organelles (Supplementary Fig. 13b). We observed a smooth transition between the ability of peptides to target mitochondrion and chloroplast as we traversed along the interpolation path. Furthermore, *in silico* analysis revealed 62 peptides exhibited a high likelihood of functioning as dual-targeting sequences (Fig. 5b; Supplementary Table 3).

**Fig. 5:**
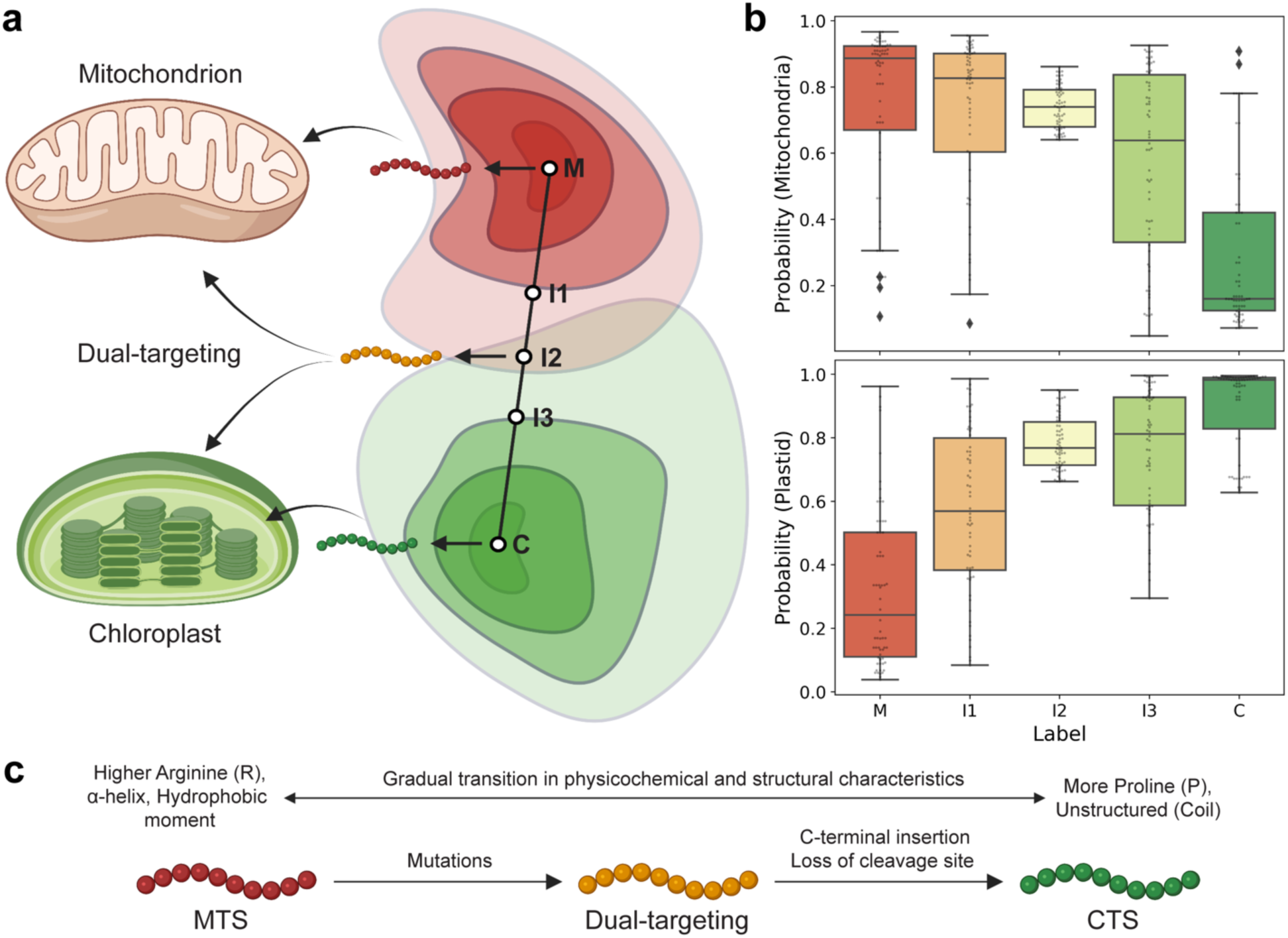
Designing dual-targeting peptides using Dual-VAE. **a)** Schematic of interpolation in the latent space to generate peptides capable of targeting both mitochondria and chloroplasts. **b)** DeepLoc 2.0 predictions for the 62 dual-targeting peptides. The likelihood of mitochondrial targeting diminishes while chloroplast targeting increases along the interpolation path from mitochondrial targeting sequences (MTS) to chloroplast targeting sequences (CTS), leading to the emergence of dual-targeting characteristics at the midpoint. **c)** Proposed evolution of dual-targeting peptides. If dual-targeting sequences evolve from MTS, they gather mutations that alter the composition of specific amino acid residues, physicochemical properties, and secondary structural features. In contrast, evolution from CTS not only requires similar mutations but also the insertion of an unstructured element at the C-terminus and modifications that incorporate a cleavage motif for recognition by mitochondrial processing peptidase.

Subsequently, we analyzed the interpolation path for these 62 putative dual-targeting peptides to understand the changes in physicochemical and structural characteristics (Extended Data Fig. 1). Examination of the amino acid composition showed a significant increase in Serine (S) and Leucine (L) and a decrease in Arginine I and Alanine (A) when compared with MTSs. An opposite trend was observed for arginine in comparison with CTSs, in agreement with a previous study that investigated the role of arginine in specificity for targeting the chloroplast^42^. CTSs were also enriched in Proline (P) when matched with MTSs and dual targeting sequences. Comparison of peptide lengths revealed a gradual increase from MTSs to interpolated sequences, while CTSs exhibited notably greater length, suggesting an insertion. Multiple sequence alignment of interpolated sequences revealed amino acid substitutions at the beginning, followed by the insertion of amino acids along the length of the peptide (Supplementary Fig. 14). Furthermore, we saw a decrease in global hydrophobic moment as the transition progressed from MTSs to CTSs, while net charge exhibited a similar distribution. Next, we employed S4PRED to predict the secondary structure of the peptides, indicating an increase in unstructured elements (coil) for dual-targeting peptides and CTSs at the expense of α-helix. These findings are consistent with a previous study that examined the evolution of MTSs and CTSs from antimicrobial peptides^43^. We also analyzed the C-terminus of these peptides and noted the conservation of the RR, R-2, or R-3 motifs in both MTSs and dual-targeting peptides (Supplementary Fig. 15), suggesting the potential for successful cleavage upon import into mitochondria. In summary, a smooth transition was observed in the physicochemical and structural characteristics while transitioning from MTSs to dual-targeting peptides. Based on these observations (Fig. 5c), we hypothesize that the dual-targeting sequences are more likely to have evolved from mitochondrial targeting sequences.

### Subcellular localization of enzymes enhances 3-hydroxypropionic acid production

As a proof of concept, we demonstrate the utility of characterized MTSs for the metabolic engineering of 3-hydroxypropionic acid (3-HP). While nature has developed various biosynthetic routes to produce 3-HP, we selected the β-alanine pathway owing to its higher theoretical yield^44^. This pathway involves three genes to convert the endogenous precursor, L-aspartate, to 3-HP: aspartate decarboxylase (PAND), β-alanine-pyruvate aminotransferase (BAPAT), and 3-hydroxypropanoate dehydrogenase (YDFG). Previously, the pathway was established in the cytoplasm of *S. cerevisiae*^44^. However, given the abundant supply of the starting precursor of this pathway in mitochondria^9^, we opted to localize the biosynthetic pathway for 3-HP in this organelle, aiming for improved production (Fig. 6a). We constructed two plasmids harboring genes without and with N-terminal MTSs (COX4-PAND, AMTS131-BAPAT, and AMTS3-YDFG), transformed them into the CEN.PK strain, cultured the strains in 2 mL of SC-URA (50 g/L glucose) media, and quantified 3-HP using HPLC. Compartmentalizing the pathway in the mitochondria led to higher 3-HP production (Fig. 6b), showing an increase of 62.3% compared to the cytoplasmic pathway (i.e., from 1.70 g/L to 2.76 g/L). Moreover, we observed 3-HP in the supernatant, indicating the presence of a putative 3-HP transporter on the mitochondrial membrane as reported elsewhere^45^, whose overexpression might further improve 3-HP titers.

**Fig. 6:**
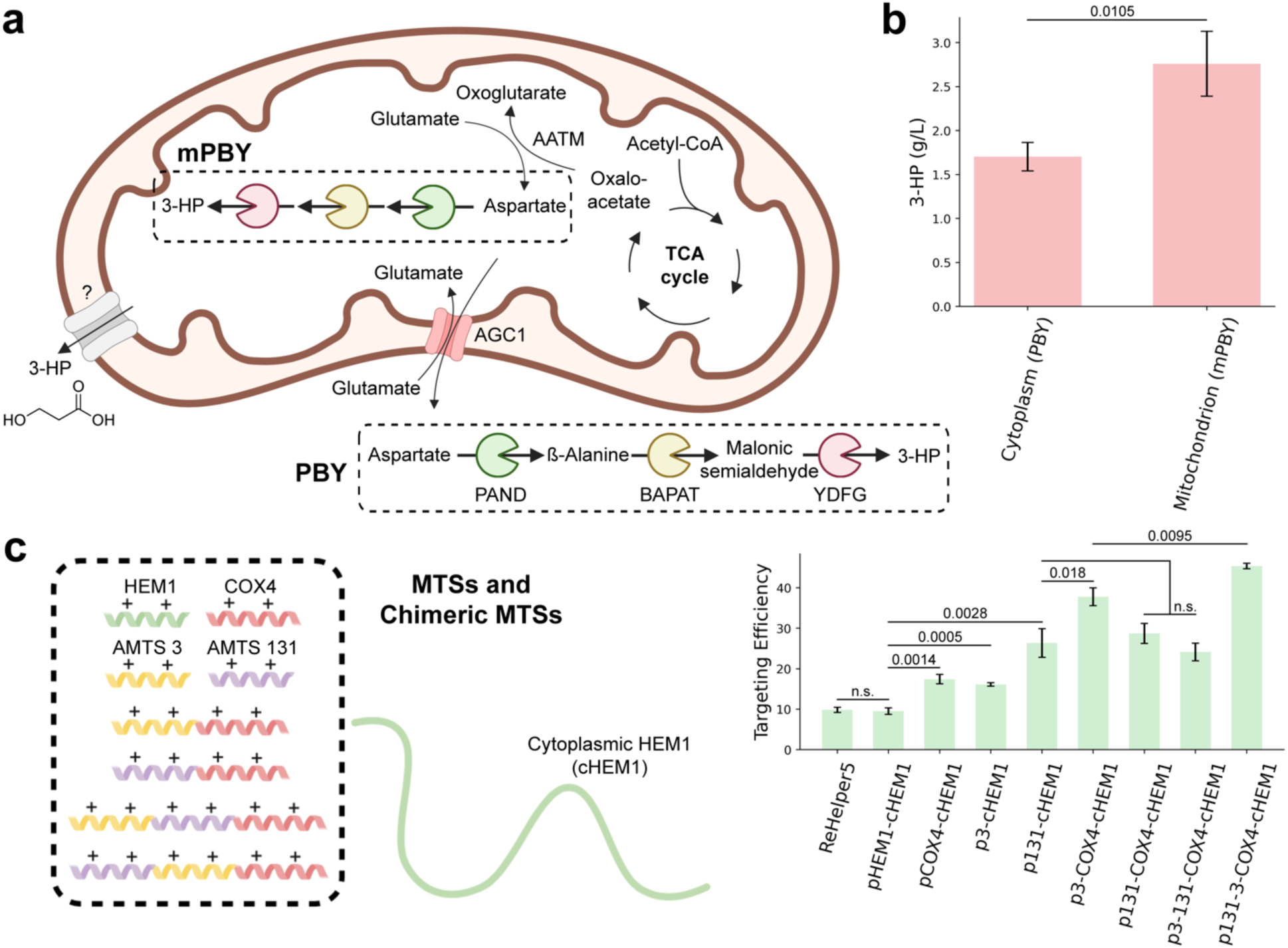
Application of characterized MTSs for metabolic engineering and protein delivery. **a)** Schematic of strains harboring plasmids for cytoplasmic (PBY) and mitochondrial (mPBY) 3-hydroxypropionic acid (3-HP) biosynthetic pathways. **b)** Production of 3-HP from SC-URA (50 g/L glucose). **c)** Enhancing mitochondrial targeting efficiency using chimeric MTSs. A library of MTSs and chimeric MTSs was cloned upstream of the HEM1 gene without the native MTS and transformed into *S. cerevisiae*. Targeting efficiency was assessed by quantifying the ratio of 5-ALA produced to mRNA levels of the HEM1 gene. PAND: aspartate decarboxylase; BAPAT: β-alanine-pyruvate aminotransferase; YDFG: 3-hydroxypropanoate dehydrogenase; HEM1: 5-aminolevulinate synthase; AATM: aspartate aminotransferase, mitochondrial; AGC1: mitochondrial aspartate-glutamate transporter. Data represents mean ± s.d. n = 3 biological replicates. *P* value was calculated by two-tailed unpaired t-test. N.s. = not significant (*p* > 0.05).

### Peptide chimeras improve mitochondrial targeting efficiency

In our second application, we showcase the application of artificial MTSs for delivering cargo proteins to mitochondria. The targeting efficiency of MTSs is dependent on the passenger protein^14^. Therefore, finding an optimal MTS requires screening a diverse library. An alternate strategy for improved targeting involves employing chimeric MTSs, i.e., a combination of MTSs in series^46,47^. To test this hypothesis, we utilized endogenous 5-aminolevulinate synthase (HEM1) in *S. cerevisiae* as the model system. HEM1 catalyzes the synthesis of 5-aminolevulinate (ALA) from succinyl-CoA and glycine within the mitochondria, the rate-limiting step in heme biosynthesis^48^. Mutations in ALAS1, a homolog of HEM1, are associated with X-linked protoporphyria and X-linked sideroblastic anemia in humans^49^. Therefore, the efficient delivery of HEM1 to mitochondria is beneficial for metabolic engineering and the development of large molecule therapeutics. However, as MTS is positioned at the N-terminus of the protein, it can also influence transcription rates and, consequently, mRNA levels. Therefore, we defined targeting efficiency as the ratio of 5-ALA produced to the mRNA levels of HEM1, a definition more relevant to targeted mRNA delivery to mitochondria. We constructed a library of chimeric MTS using COX4, AMTS3, and AMTS131. For higher-order combinations, we maintained COX4 at the C-terminus to ensure cleavage and no interference with downstream protein folding.

We introduced these MTS/chimeric MTS-cHEM1 constructs into *S. cerevisiae* and subsequently measured both 5-ALA production and mRNA levels of the HEM1 gene to evaluate targeting efficiency (Fig. 6c and Supplementary Fig. 16). For individual targeting sequence, COX4 and AMTSs exhibited a significant increase in targeting efficiency compared to the native MTS. Furthermore, we observed that the targeting efficiency increased with the number of MTSs in chimeric constructs. Specifically, a 4.76 and 2.53-fold enhancement was achieved with three MTSs in series, emphasizing the importance of the number and order of MTSs for mitochondrial targeting. However, 5-ALA production in chimeric constructs did not show significant improvement over the individual MTSs, primarily due to lower mRNA levels, which may be attributed to suboptimal codon optimization.

## Discussion

Mitochondria play a key role in energy metabolism, biosynthesis of macromolecular precursors and reducing equivalents, and effective management of metabolic waste. Therefore, mitochondrial targeting holds immense promise for metabolic engineering and therapeutics. However, despite the crucial role of mitochondria in cellular metabolism, the repertoire of well-characterized mitochondrial targeting sequences (MTSs) remains limited. This scarcity restricts the options available for directing proteins to mitochondria, potentially leading to suboptimal targeting efficiency and compromised functionality of the delivered cargo or pathways. These limitations underscore the urgent need for the design and characterization of diverse, functional MTSs.

In this study, we addressed this challenge by developing a deep generative AI framework to design artificial mitochondrial targeting sequences. First, we curated a large dataset of MTSs occurring in nature and trained a Variational Autoencoder on these peptide sequences. We validated the functionality of the generated MTSs using DeepLoc 2.0. A detailed analysis of physicochemical and structural attributes revealed the sequences were positively charged, amphiphilic, and tended to form an α-helix, features important for targeting mitochondrion. Next, we devised a sampling scheme to prioritize MTSs for experimental validation. Using confocal microscopy, we characterized 41 new-to-nature peptides *in vivo* in four eukaryotic species, achieving a success rate of 50-100%. Moreover, we demonstrated that the peptides are successfully cleaved *in vivo* in the HEK293 cell line.

Furthermore, our analysis showcased that some of the generated sequences are likely capable of targeting both the mitochondrion and chloroplast. Therefore, we trained another model, Dual-VAE, on MTSs and CTSs from the Viridiplantae kingdom. Subsequently, we utilized linear interpolation in the latent space and generated 62 putative dual-targeting sequences. We analyzed the variations in the features of targeting sequences along the interpolation trajectory and provided insights into how dual-targeting sequences may have likely evolved from mitochondrial targeting peptides. In the future, we anticipate that the characterization of these peptides will enhance our understanding of dual-targeting in plants and boost titers of biochemicals, such as taxenes^50^, by utilizing acetyl-CoA from both compartments. Lastly, we demonstrated the application of the characterized peptides for metabolic engineering and protein delivery. We localized the enzymes in the β-alanine pathway to produce 3-HP in the mitochondria and observed a 1.62-fold improvement in titers compared to the cytoplasmic counterpart. In the second application, we focused on improving the targeting of HEM1 to mitochondria using chimeric MTSs and noticed a 4.76-fold enhancement in targeting efficiency. We saw the order and number of MTSs in a series combination are both important to enhance localization.

A promising avenue for further improvement lies within the model architecture. One could directly incorporate the organism label and protein information using a Conditional Variational Autoencoder^51^ or frame the problem as a machine translation task^19^, removing the need for selective sampling. In addition, it would capture the bias introduced through interaction with the import machinery and the intrinsic relation between the passenger protein and MTS. One could also streamline the selection of artificial MTSs for *in vivo* characterization by inputting vectors positioned close to the cluster center (derived from VAE latent vectors of MTSs in a proteome) into the decoder, facilitating the generation of artificial MTS tailored to the specific organism under study.

Moreover, developing a high-throughput assay would significantly improve efforts in characterizing artificial MTSs designed in this study. Here, we utilize fluorescent protein tagging and subsequent *in vivo* microscopy, a technique fundamental for studying protein localization in cellular biology^52^. However, this method is not suitable for quantifying the targeting efficiencies at scale. Therefore, deploying quantitative assays based on the self-assembling split green fluorescent protein (Split-GFP) technology^53,54^, fluorescence-activated cell sorting (FACS), and next-generation sequencing will generate high-quality sequence-to-targeting efficiency data and aid in identifying the best MTS for the passenger protein of interest. Furthermore, one could utilize the dataset to train a supervised ML model and screen MTSs or chimeric MTSs *in silico*. However, based on our HEM1 delivery application, it is evident that swapping MTSs at the N-terminus significantly affects RNA levels. Therefore, it would be more logical to use codon language models^55,56^ to represent MTSs, capturing both transcriptional and targeting efficiencies.

In conclusion, our VAE models are highly capable of designing diverse, functional mitochondrial and new-to-nature dual-targeting sequences, improving our ability to deliver necessary cargo for metabolic engineering and biomedical applications.

## Methods

### Variational Autoencoder

To train the VAEs, targeting peptides from the curated MTS dataset were one-hot encoded. Encoders and decoders of fully connected layers were trained using the loss function (Eq. 1), consisting of the reconstruction loss and Kullback−Leibler (KL) divergence^57^. The reconstruction loss was calculated using the binary cross-entropy function as the difference between the reconstructed output and the input data, encouraging the VAE to produce reconstructions that closely resemble the original data, while the KL divergence term was minimized to regularize the latent space by penalizing deviations from a standard normal distribution. The training process was stopped when the validation loss did not improve for five consecutive epochs. Hyperparameters, including the annealing rate, were optimized for both models.

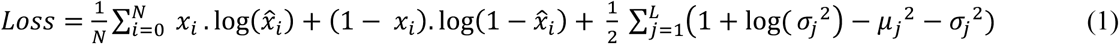

where *x_i_* and *x^^^_i_* are ground truth and reconstructed output (of size *N*) at the position I and *μ*_*j*_, σ_*j*_, and *L* are mean, variance, and dimensionality of the latent variable **z**, respectively.

### Sampling and annotating sequences for *in vivo* validation

Artificial MTSs were generated by feeding random numbers drawn from a normal distribution N(μ = 0, σ = 1) as inputs to the decoder. The resulting reconstructions were converted into amino acid sequences employing the dictionary used for one-hot encoding. Sequences preceding the cleavage symbol ‘$’ were extracted and subjected to validation checks. Subsequently, these sequences were assigned organism labels based on their proximity to the cluster center (Eq. 2). Moreover, consistency in the local space was ensured by assessing which organism most of the 20 nearest neighbors belong to, verifying their alignment with the assigned label. Both artificial and naturally occurring MTSs were encoded using the pre-trained UniRep model ***U***. The cluster center ***C**_o_* was computed as the mean of UniRep embeddings for MTSs within an organism O’s proteome (Eq. 3), while the nearest neighbors were identified by measuring the Euclidean distance between the UniRep embeddings of artificial peptides and MTSs of all four distinct eukaryotic organisms (Eq. 4). Finally, artificial MTSs were ranked according to their distances from the cluster centers of individual organisms and the top eight sequences were chosen for subsequent *in vivo* validation.

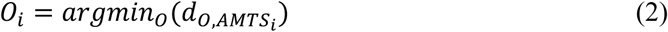

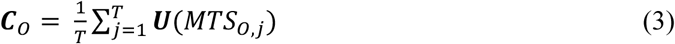

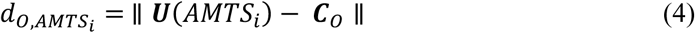

where T is the total number of MTSs in the proteome of organism O, and O is one of the following: *S. cerevisiae*, *R. torulodies*, *N. tabacum*, or *H. sapiens*.

### Analysis of artificial targeting sequences

All VAE-generated sequences were fused to the N-terminal of GFP (without the start amino acid, methionine) and analyzed for functionality using the local software of DeepLoc 2.0^4^. The generated output was analyzed in-house based on the predicted subcellular localization or provided probability thresholds. Physicochemical features, including amino acid composition, net charge, Eisenberg hydrophobicity, and GRAVY score, were calculated using Biopython, while the hydrophobic moment was obtained using modlAMP^30^. Secondary structures of the artificial peptides were predicted using the S4PRED model^32^. Sequence logos for the analysis of cleavage sites were generated using WebLogo^58^. Clustal Omega^59^ and pyMSAviz were used to create and visualize MSAs, respectively.

### Profile Hidden Markov Model

The HMMER package^27^ was employed with default settings to design MTSs with the pHMM. A multiple sequence alignment was obtained for 4,000 MTSs reported in Swiss-Prot using Clustal Omega^59^ and was fed to HMMER’s hmmbuild to create an HMM profile. Subsequently, HMMER’s hmmemit was utilized to generate 1000 putative MTSs, from which 730 sequences were randomly sampled and fused to GFP at the N-terminus to assess functionality using DeepLoc 2.0.

### Strains, media, and reagents

*E. coli* strain NEB10β (New England Biolabs, MA) was used for all cloning experiments. The following reference strains were used in the study: *S. cerevisiae* BY4741 (*MATα his3Δ1 leu2Δ0 met15Δ0 ura3Δ0*), *S. cerevisiae* CEN.PK, *R. toruloides* 880CF (IFO0880-pANT1-SpCas9-pGAPDH1-MaFAR), *Agrobacterium tumefaciens* strain GV3101 (pMP90), tobacco *Nicotiana benthamiana*, and HEK293T (ATCC #CRL-3216). Reagents, buffer components, and growth media were purchased from Millipore Sigma (Burlington, MA), Qiagen (Germany), or Thermo Fisher Scientific (Hampton, NH). Restriction enzymes, T4 polynucleotide kinase, T4 DNA ligase, Q5 DNA polymerase, NEBuffer 3.1, Gibson Assembly Master Mix, and HiFi DNA Assembly Master Mix were purchased from New England Biolabs, MA. PrimeSTAR Max DNA Polymerase was purchased from Takara Bio. WT *Pf*Ago and *Pf*Ago* aliquots were obtained as previously described^35^. The plasmid CD3-991 (35S::ScCOX4-mCherry) was procured from the Arabidopsis Biological Resource Center (ABRC). All DNA oligonucleotides were ordered from Integrated DNA Technologies (IDT) (Coralville, IA), while synthetic genes were codon-optimized for *S. cerevisiae* using IDT’s Codon Optimization Tool and purchased from Twist Biosciences (San Francisco, CA). Targeting sequences were codon-optimized using IDT’s Codon Optimization Tool or Juggler tool in BOOST^61^ and ordered as oligos or gBlocks. Oligos, primers, and synthetic genes used in this study are listed in Supplementary Table 4.

### Plasmid construction

Scarless DNA assembly for the artificial MTSs was performed using *Pf*Ago and T4 DNA ligase. Guide DNA phosphorylation, *Pf*Ago/AREs cleavage of backbone fragments, and purification of *Pf*Ago cleaved products were carried out as described elsewhere^35^. Plasmid backbones were amplified using PrimeSTAR Max DNA Polymerase and digested with WT *Pf*Ago and *Pf*Ago*. To prepare the inserts, oligos containing the MTSs with protruding sticky ends were phosphorylated using T4 polynucleotide kinase. The reaction mixture was incubated at 37 °C for 1 h, 65 °C for 20 min, and cooled to 4 °C. An equimolar mixture of phosphorylated oligos was annealed in NEBuffer 3.1. The mixture was heated to 94 °C for 3 min and gradually cooled to 25 °C over a period of 45 min. The annealed product and the digested backbone were assembled using T4 DNA ligase. Subsequently, 3 µL of the ligation mixture was added to 50 µL of chemically competent NEB10β cells and transformation was performed following the manufacturer’s protocol. The initial library of VAE-generated MTSs tested in *S. cerevisiae* was cloned using Gibson Assembly^33^. To create an NLS construct, pUC19_SV40_EGFP was cut with *Bsa*AI and then assembled using HiFi DNA assembly.

Correct assembly was first verified using Colony PCR and finally confirmed by Sanger sequencing. For plasmid amplification, the colony with the correct plasmid was cultured in LB medium supplemented with appropriate antibiotics. The plasmid DNA was purified from the cultures using the QIAprep Spin Miniprep Kit (Qiagen, Germany), following the manufacturer’s protocol. The plasmids constructed in this study are listed in Supplementary Table 5.

### Analysis of MTS-GFP localization by fluorescence microscopy

To characterize MTSs in *S. cerevisiae*, AMTS-EGFP plasmids were transformed into *S. cerevisiae* using the commonly used lithium acetate heat shock protocol^62^. Cells with the AMTS-EGFP plasmids were inoculated overnight into 2 mL SC-URA media and cultivated at 30 °C and 250 rpm for approximately 18 h. When OD600 reached 0.1–0.3, cells were centrifuged, and the media was replaced with 1 mL Phosphate-Buffered Saline (PBS; without calcium and magnesium) containing 200 nM of MitoTracker^®^ Orange CMTMRos (purchased from Invitrogen, USA and diluted in dimethyl sulfoxide to create 1 mM stock solution). The cells were stained at 30 °C for 15 min (culture tube was covered with foil, 250 rpm), pelleted, washed with 1 mL PBS twice, resuspended in PBS with 2% glucose, and transferred to a 1.5 mL microcentrifuge tube. The stained yeast cells were mounted onto Poly-L-Lysine coated slides (Cole-Parmer) and analyzed using the ZEISS LSM880 Confocal Laser Scanning Microscope. Imaging was performed using a 63x oil immersion objective lens. A similar procedure was followed for *R. toruloides* strains expressing the integrated AMTS-EGFP construct with minor modifications. Cells were cultivated in SC (pH 5.6) and stained with Rhodamine B, hexyl ester (Yeast Mitochondrial Stain Sampler Kit purchased from ThermoFisher Scientific).

To investigate MTSs in plant cells, *Agrobacterium tumefaciens* GV3101 (pMP90) transformed with the plasmids AMTS-YFP and CD3-991 were cultured overnight in 2 mL LB medium containing appropriate antibiotics at 28 °C. Bacterial cells were collected at 10,000 rpm for 1 min and resuspended with 2 mL water to wash out the antibiotics, followed by one more washing. Cells were then resuspended in the infiltration buffer (10 mM MgCl_2_, 10 mM MES pH 5.7, 200 μM acetosyringone) and the concentration was adjusted to OD600 = 0.6. Cells in the buffer were placed at room temperature for two hours before infiltration. Agrobacterium solution was infiltrated into leaves through a syringe without a needle. After three days, the fluorescence on the epidermis was observed and images were captured on a ZEISS LSM 710 confocal microscope. YFP was excited at 514 nm and emission was collected at 520-540 nm, while mCherry was excited at 561 nm and emission was collected from 580 to 620 nm.

HEK293T cells were transfected with 200 ng of MTS-GFP plasmids using Fugene HD (Promega #E2311) when the cell confluence reached approximately 60%. Transfection was carried out in a 12-well plate on Poly-L-lysine-coated glass coverslips. The growth media was changed at 24 hours and 48 hours. Post-transfection, live cells were incubated with MitoTracker^®^ Orange (Thermo Fisher #7510) for 30 minutes. The cells were fixed in 3.7% Formaldehyde for 15 minutes, followed by three 5-minute washes with 1xPBS. Subsequently, the coverslips were mounted on ProLong Diamond with DAPI (Thermo Fisher # P36962). The images were acquired at 100x magnification on an OMX-V4 microscope (GE Healthcare) equipped with a U Plan S-Apo 100x/1.40-NA oil-immersion objective (Olympus). The images were deconvolved using the previously described deconvolution parameters^63^ and final images were created using Fiji^64^.

### Confirmation of MTS cleavage in HEK293 cells using Western Blot analysis

HEK293 cells were transfected with 500 ng of purified plasmid DNA using Lipofectamine 3000 (Invitrogen) in a 12-well plate for 48 hours. Subsequently, the cells were lysed in 150 µl of 1x sample buffer (Bio-Rad #1610747) containing 5% β-mercaptoethanol and heated at 95 °C for 5 minutes. For the GFP western blot, the cell lysates were normalized to ensure comparable GFP levels and were then resolved on a 14% SDS-PAGE gel. The proteins were transferred onto a PVDF membrane (Bio-Rad #1704156) and incubated overnight with primary antibodies (GFP, Thermo Fisher #MA1-952, 1:2500; β-Tubulin, Proteintech #66240, 1:20,000) in 5% non-fat milk. Afterward, the membrane was incubated for 20 minutes with an HRP-conjugated goat anti-mouse secondary antibody (Jackson ImmunoResearch #115-036-003). The membrane was developed using a western ECL substrate (Bio-Rad #1705061) and imaged on Amersham ImageQuant studio.

### Strain construction

Prior to the random integration of the AMTS-EGFP expression cassette, the deletion of the carotenogenic reporter gene *CAR2* was performed. *R. toruloides* 880CF was transformed with the gRNA expression cassette along with the *hpt* gene encoding for hygromycin phosphotransferase amplified from a previously constructed pRTH-car2d vector^37^. A white colony demonstrating the phenotype of a successful knockout was picked for subsequent experiments. 1000 ng of AMTS-EGFP expression cassette along with the *nat* gene conferring the nourseothricin resistance was amplified using PCR, transformed into *R. toruloides* 880CF-*CAR2*Δ strain, and characterized using fluorescence microscopy. Transformation of *R. toruloides* was performed using the modified lithium acetate protocol as described previously^37^. The strains constructed in this study are listed in Supplementary Table 6.

### Quantification of 3-HP in engineered strains

The 3-HP producing strain, SCE-R2-200, was obtained from a previous study^44^. PAND, BAPAT, and YDFG genes were amplified from this strain, while promoters and terminators were sourced from the genome of the CEN.PK strain. Mitochondrial targeting sequences (COX4, AMTS131, AMTS3) were amplified from the MTS-GFP plasmids. All fragments included 60 bp overlaps with adjacent fragments, and plasmid construction with pTH backbone was carried out using DNA assembler^65^. The constructed plasmids were then transformed into the CEN.PK strain via the commonly used lithium acetate heat shock protocol^62^. Three colonies for each strain were picked from the SC-URA plate and inoculated in a culture tube containing 2 mL of SC-URA media (50 g/L glucose). After five days of incubation at 30 °C and 250 rpm, the cultures were centrifuged, and the supernatant was collected and diluted twofold. The samples were then directly subjected to HPLC analysis. HPLC was performed using an Agilent 1260 system (Agilent, USA) with an Aminex HPX-87H Column #1250140 (BioRad, USA). The column was operated at 60 °C with 2.5 mM H_2_SO_4_ at a flow rate of 0.6 mL/min. The 3-HP peak was observed at 12.2 minutes.

### Measuring targeting efficiency of chimeric MTSs

For 5-ALA quantification, 200 µL of culture medium was collected and mixed with 200 µL of 1 M acetate buffer (pH 4.6) and 100 µL of acetylacetone. The mixture was heated at 100 °C for 10 min. Subsequently, the mixture was diluted tenfold and mixed with an equal volume of modified Ehrlich’s reagent (0.1 g/mL p-dimethylaminobenzaldehyde (DMAB) and 0.16 mg/mL hypochlorous acid in glacial acetic acid) and kept at room temperature for 10 min. The formation of a pink compound was observed and the absorbance was measured at 553 nm using a SpectraMax Mini Multi-mode microplate reader (Molecular Devices, USA). For qPCR, strains harboring the HEM1-expressing plasmid were cultured in SC-URA for one day, and then total RNA was extracted using the Rneasy Mini Kit (Qiagen, USA). The total RNA was subjected to reverse transcription to obtain cDNA using the iScript™ Reverse Transcription Supermix (BioRad, USA). qPCR was performed using iTaq Universal SYBR Green Supermix (BioRad, USA) in the QuantStudio™ 6 Pro Real-Time PCR System (Thermo, USA). The HEM1 qPCR primers used were as follows: Forward: CAGGAGTCGGGTTTCGATTAC, Reverse: CTTGGCCAATCGGTTGATATTG. RNA extraction, cDNA synthesis, and qPCR were performed according to the manufacturer’s protocol.

## Data availability

The data supporting the findings of this study are available within the article and its Supplementary Information files or uploaded through public repositories. If specific data is believed to be missing, that data is available from the corresponding author upon request.

## Code availability

The source code and the trained VAE models developed in the study are available on GitHub at: https://github.com/Zhao-Group/MTS-VAE. Datasets curated for training the VAE models and analysis are on Zenodo: https://zenodo.org/records/13214860. The scripts for generative models and data analysis were built on Python 3.9.16, PyTorch 1.9.1, Numpy 1.24.3, Scipy 1.10.1, BioPython 1.81, Scikit-learn 1.1.3, Pandas 1.5.3, modlAMP 4.3.0, and CD-HIT 4.8.1.

## Acknowledgements

This work was funded by the DOE Center for Advanced Bioenergy and Bioproducts Innovation (U.S. Department of Energy, Office of Science, Biological and Environmental Research Program under Award Number DE-SC0018420). Any opinions, findings, and conclusions or recommendations expressed in this publication are those of the author(s) and do not necessarily reflect the views of the U.S. Department of Energy. The online tool BioRender (biorender.com) was used to create Fig. 1, Fig. 3, Fig. 5a, Fig. 6, and Supplementary Fig. 12. We thank Prof. Irina Borodina (DTU Biosustain) for providing the 3-HP producing strain, SCE-R2-200. We thank Dr. Behnam Enghiad for his suggestions on *Pf*Ago-based assembly and Dr. Austin Cyphersmith from Core Facilities at the Carl R. Woese Institute for Genomic Biology for his help with the fluorescence microscopy. The authors acknowledge the use of computing facilities of Biocluster at the Carl R. Woese Institute for Genomic Biology and Nano cluster at the National Center for Supercomputing Applications. This work also used the Delta system at the National Center for Supercomputing Applications through allocation BIO230077 from the Advanced Cyberinfrastructure Coordination Ecosystem: Services & Support (ACCESS) program, which is supported by National Science Foundation grants #2138259, #2138286, #2138307, #2137603, and #2138296.

## Contributions

A.G.B. and H.Z. conceived and designed the study. A.G.B. performed the computational experiments. A.G.B., S.-I.T., A.Z., N.S., X.X., S. Z., and T.A.M. performed the wet-lab experiments, and analyzed the data. A.G.B., S.-I.T., N.S., X.X., L.-Q.C., and H.Z. wrote the manuscript.

## Competing interests

The authors declare no competing interests.

---

**Extended Fig. 1:**
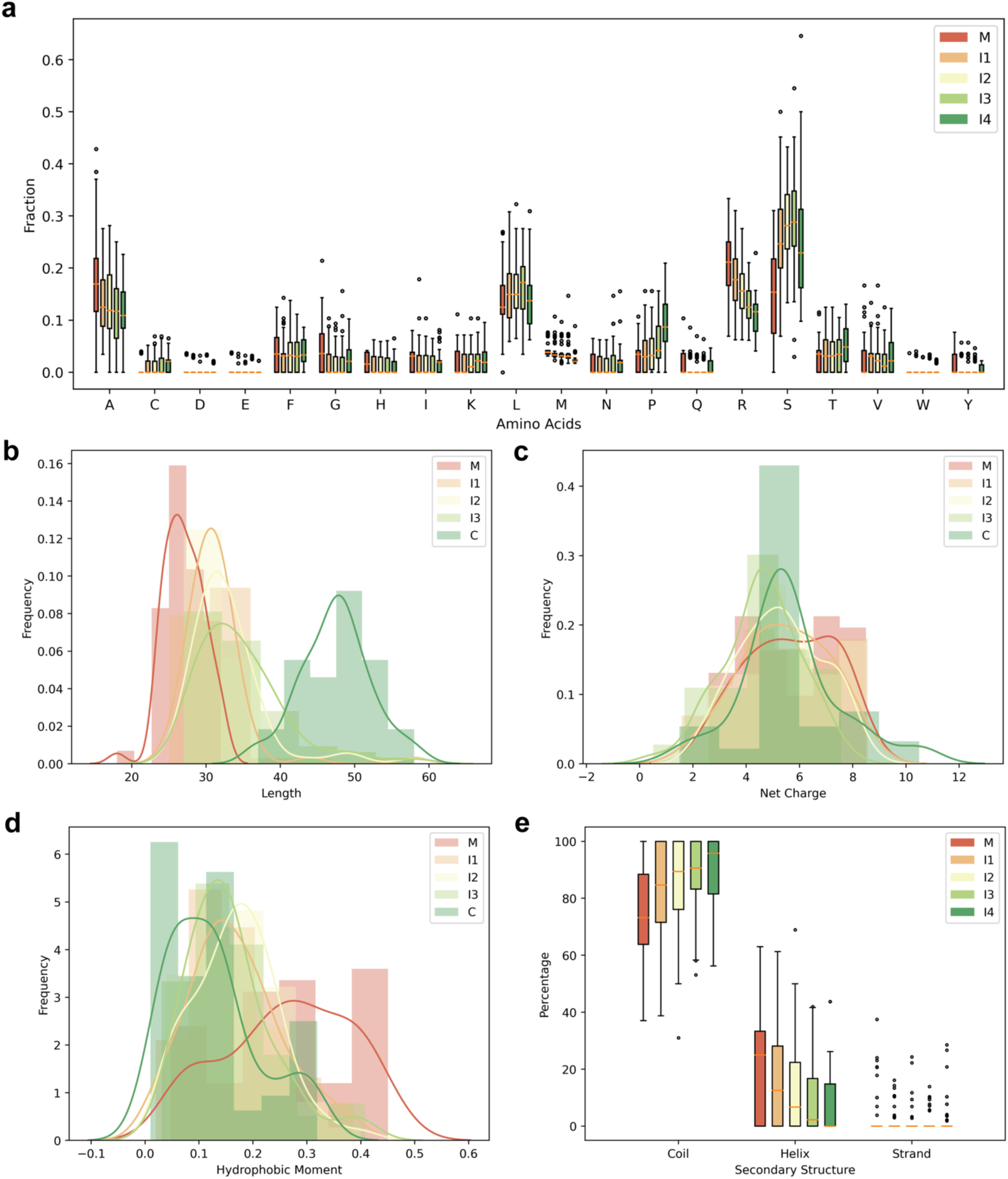
Physicochemical and structural features of artificial sequences sampled along the interpolation path of dual targeting peptides, including their **a)** Amino acid composition, **b)** Length, **c)** Net charge, **d)** Hydrophobic moment, and **e)** Secondary structure. A smooth transition is observed for various attributes, such as amino acid composition (arginine, proline), hydrophobic moment, and secondary structure elements when transitioning from mitochondrial to chloroplast targeting sequences.

## Notes

### Competing Interest Statement

The authors have declared no competing interest.

